# A comprehensive assessment of benign genetic variability for neurodegenerative disorders

**DOI:** 10.1101/270686

**Authors:** Rita Guerreiro, Celeste Sassi, Jesse Raphael Gibbs, Connor Edsall, Dena Hernandez, Kristelle Brown, Michelle K Lupton, Laura Parkinnen, Olaf Ansorge, Angela Hodges, Mina Ryten, for the UK Brain Expression Consortium, Pentti J. Tienari, Vivanna M. Van Deerlin, John Q Trojanowski, Kevin Morgan, John Powell, Andrew Singleton, John Hardy, Jose Bras

## Abstract

Over the last few years, as more and more sequencing studies have been performed, it has become apparent that the identification of pathogenic mutations is, more often than not, a complex issue. Here, with a focus on neurodegenerative diseases, we have performed a survey of coding genetic variability that is unlikely to be pathogenic.

We have performed whole-exome sequencing in 478 samples derived from several brain banks in the United Kingdom and the United States of America. Samples were included when subjects were, at death, over 60 years of age, had no signs of neurological disease and were subjected to a neuropathological examination, which revealed no evidence of neurodegeneration. This information will be valuable to studies of genetic variability as a causal factor for neurodegenerative syndromes. We envisage it will be particularly relevant for diagnostic laboratories as a filter step to the results being produced by either genome-wide or gene-panel sequencing. We have made this data publicly available at www.alzforum.org/exomes/hex.

## Introduction

One of the most important objectives in medical research is the complete understanding of how genetic variability influences a specific phenotype or trait. Methods to accomplish this at a genomic level have become available over the last decade with the introduction of genome-wide genotyping and “next-generation” sequencing (Bras et al., 2012).

The widespread adoption and application of sequencing technologies allows two main types of studies: 1) sequencing of large cohorts of cases and controls to perform association studies with virtually complete genomic information; and 2) sequencing of individuals from families where a specific disease segregates. If the first application requires the analysis of as many individuals as possible, the second one requires instead a strict filtering of variants to allow for the identification of the causative one(s) (Goldstein et al., 2013).

As sequencing technologies have matured over the last few years, the main issue researchers have faced has changed from dealing with false positive findings to dealing with the sheer number of true positive results. Because of the large number of rare, and sometimes private variants, it is often difficult to determine if a specific variant is benign in nature or if, instead, it is pathogenic (Biesecker et al., 2012).

The most common approach to filter variants has been to use publicly available databases of genetic data, such as the 1000Genomes (Abecasis et al., 2010), the Exome Aggregation Consortium (ExAC; URL: http://exac.broadinstitute.org/) or more recently the genome Aggregation Database (gnomAD: http://gnomad.broadinstitute.org/) because these resources provide users with genetic variability data on large numbers of individuals (Lek et al. 2016).

Although this type of databases is very powerful, it should be noted that they do not provide detailed phenotypic information on the subjects included. For example, although cohort-level disease information is sometimes included, they do not provide individual-level information (such as the ages of the participants). This is essential information for studies that deal with diseases that occur later in life, as is the case for many neurodegenerative diseases. It is not clear, for example, how many samples have developed dementia since their inclusion in these studies/cohorts. As such, using these data to naively filter variants from cases presenting with any form of dementia can bias the results.

Consequently, there is a need for a resource that allows researchers to confidently filter out variants from their own family-based sequencing studies, without the risk of excluding potentially disease-causing mutations. To address this issue, and focusing on neurodegenerative disorders, we are performing whole-exome sequencing in brain tissue samples from individuals who lived to old age and had no signs of overt neurodegeneration on neuropathological examination (Braak stage <= II).

Here, we present results from the first public release of this dataset and describe the resource where the data is made publicly available (www.alzforum.org/exomes/hex).

## Results

The data present herein is based on high-coverage exome-sequencing data from 478 elderly individuals who presented no signs of neurodegeneration when subjected to a post-mortem neuropathological examination. We have generated an average of 9Gb of data per individual, which translates into a mean coverage of the exome portion of the genome of 57x and leading to the identification of a total of 1,145,559 high-quality variants with 276,997 of these variants directly affecting the coding sequence.

### Variants in known neurodegenerative disease genes

One of the most important aspects of this resource is that it enables researchers to identify rare genetic variability in known disease-causing genes and immediately be aware that that variability will not account for a neurodegenerative disorder.

Of the variability identified in known neurodegenerative disease genes a few should be highlighted: two novel variants in *PSEN2*, nine novel variants in *LRRK2* and the variants p.Val476Ala and p.Thr763Ala in VPS35, for which reported minor allele frequencies are very low in ExAC (4.118e-05 and 8.236e-06, respectively). Each of these variants, if found in affected cases with the corresponding diseases, could be classified as disease causing, but the fact that we see them in HEX strongly argues against that interpretation. In addition to novel and low frequency variants, we have also identified variants that are reported to be pathogenic: the p. Ser163Leu and the p.Asp472Ala in *PSEN2* are both seen in our cohort, as is p.Arg1628Pro in *LRRK2*, implying they are not fully penetrant disease-causing mutations in the Caucasian population.

### Genes with loss-of-function variants

Similarly, it is also of great importance to understand, for neurodegenerative diseases, how many and which genes can be tolerated to have either one or two loss-of-function (LOF) alleles. We have identified 15,829 LOF sites, with 1,442 being LOF homozygous in at least 1 sample. Looking at single nucleotide variants specifically, given the higher error rates in indel calling, there are 3,472 nonsense variants with 273 being homozygous in at least one sample; 160 stop-loss, of which 32 are homozygous in at least one sample. Among the single nucleotide variants that are predicted as loss of function, 1,716 are novel as of dbSNP142. These variants represent a background of data where we know loss of function is unlikely to give rise to neurodegenerative disease.

### Variants previously described to be disease causing

To search for variants that have been reported as disease causing for neurological syndromes, we used dbSNP’s variant annotation of pathogenicity based on the 30/10/2017 build of the clinvar file (ftp://ftp.ncbi.nlm.nih.gov/pub/clinvar/vcf_GRCh37). A total of 696 individual variants from our cohort are annotated in dbSNP as being pathogenic.

As selected examples, and in addition to the two *PSEN2* and *LRRK2* mutations mentioned above, we identified one sample presenting a known pathogenic mutation in *SCN9A* (p.Asn642Tyr; rs121908918). This gene is responsible for a range of neurological conditions (OMIM:603415), and this particular mutation was associated with a broad clinical spectrum including generalized epilepsy and febrile seizures (Singh et al., 2009). A previously reported pathogenic mutation in *SNCB* was also identified in this cohort (p.Pro123His; rs104893937). This mutation was reported to cause dominant Lewy Body dementia (Ohtake et al., 2004). Three individuals (ages at death ranging from 73 to 83 years) presented with the p.Arg468His mutation (rs138382758) in MFN2, previously described to cause Charcot-Marie-Tooth disease in a dominant fashion (Casasnovas et al., 2010). Finally, we identified a single case presenting the p.Pro80Leu variant in *CHCHD10* a known cause of FTLD/ALS in an individual with age at death over 80 years.

### Public release of data as HEX (Healthy EXomes database)

A resource such as this has tremendous potential in enabling researchers and molecular diagnosticians to perform much more complete and confident filtering of genetic variants. We have made this data publicly available as part of Alzforum at www.alzforum.org/exomes/hex. This offers the added benefit of allowing the user to integrate different types of information from a variety of sources and databases, without leaving the same website.

## Discussion

Over the past few years we have been committed to study benign genetic variability in genes known to cause dementia in order to better understand the pathogenic role of each variant in these genes. To do this, we have systematically reassessed the pathogenicity of variants described in the genes known to more frequently harbor Alzheimer’s disease and frontotemporal dementia causative mutations (*APP*, PSEN1, *PSEN2*, *GRN* and *MAPT*) (Guerreiro et al., 2010a; Guerreiro et al., 2010b). To follow up that work we have now performed exome sequencing in 478 samples from elderly individuals who presented no signs of neurodegeneration on post-mortem neuropathological examination. Given that we now know that plaque and tangle pathology occur 10-20 years prior to clinical onset of disease, even the younger individuals in our cohort would be unlikely to develop disease for at least 15-20 years (Weiner MW et al., 2017). Because of the sample selection that was performed, each of the variants present in this cohort is highly unlikely to be a penetrant causal mutation for a neurodegenerative condition.

It is often difficult to establish pathogenicity for novel genetic variants and several examples of miss-assignment of pathogenicity to benign variants can be found in the literature. The most common example is to assign pathogenicity to variants simply because they are found in genes that have been previously associated with a particular disease. This can be problematic from a clinical genetics perspective because it can lead to incorrect information being given to patients and family members; but it is also problematic from a fundamentally scientific perspective, because it has the potential to mislead research on the basic mechanisms underlying pathological processes. As an example, the *MAPT* variant p. Ala239Thr (rs63750096), found in two of HEX samples, was initially observed in an FTD patient with atypical brain pathology (Pickering-Brown et al., 2002). This variant was considered to be pathogenic until *GRN* was found to be associated with FTD and subsequent sequencing of *GRN* in this case revealed the presence of a partial deletion of exon 11 (Pickering-Brown et al., 2006). The finding of this probable null mutation in *GRN* in combination with FTD tau negative pathology clearly pointed to a pathogenic role of the *GRN* mutation and a benign nature of the *MAPT* variant.

It should be noted that no validation of individual variants was performed for any of the variants identified. It is thus possible that some of the variants we identify are false-positives. Some of these may be derived from regions of low sequencing coverage, while others will be technology dependent errors. The latter will also be a useful resource for researchers working on the same type of data, as identifying false positives is not always easy, particularly when one does not have access to large numbers of samples sequenced using the same technology.

The main objective of the HEX database is not to perform extensive validation of which variants may or may not be disease causing, instead we aim to provide a resource of genetic variability, created using well established analytical approaches, that was generated from very well characterized control samples for neurodegenerative diseases. The identification of variants of unknown pathogenicity, or even ones previously reported to be pathogenic, will be one of the major aspects of HEX, but also identifying genes that can tolerate loss-of-function variants and amino-acid positions that can be variable in the genome, with no effect on causing a neurodegenerative condition.

It should be noted that low frequency, mid to high-risk variants are expected to be seen in HEX. Examples of this would be the p.Asn409Ser variant in *GBA*, or the p.Arg47His variant in *TREM2*, since neither of them is disease-causing in a dominant manner and, much like the epsilon4 allele of APOE in Alzheimer’s disease, having these variants alone is neither sufficient nor necessary for the development of Parkinson’s or Alzheimer’s diseases. Additionally, it is also plausible that causative variants with incomplete penetrance are present in the database, and this should be taken into account if the mode of inheritance of the samples being studied suggests incomplete penetrance. Thus, risk variants, both high and low risk, are expected to be seen in HEX, albeit with a lower frequency than in the general population.

In summary, we describe a resource of genetic data, derived from samples of individuals who were of old age at time of death and who had no signs of neurodegeneration on neuropathological examination, and we make these data publicly available. We fully expect to continue to grow this database by including more samples and updating analytical tools as better methods are developed.

## Materials and Methods

### Sample selection

We have currently included 478 samples derived from several UK and US brain banks. As inclusion criteria we used the following: 1) age at death > 60 years; 2) no clinical symptoms of neurodegenerative disease; 3) normal neuropathological examination (since some samples were derived from individuals > 85 years, we have included samples with Braak staging of up to II in some cases); 4) unrelated individuals.

### Sample preparation and sequencing

DNA was extracted from frozen brain homogenates according to standard practices. Libraries were prepared according to Illumina’s TruSeq recommendations and sequenced on the HiSeq2000 on 2×100 base pair reads.

### Data analysis

Raw fastq files were processed according to GATK’s Best Practices. In short, reads were aligned using bwa (Li and Durbin, 2009), optical duplicates were excluded using picard (http://picard.sourceforge.net/), local realignment around indels and base quality score recalibration were performed using GATK v3.4 (Depristo et al., 2011; McKenna et al., 2010). Variant calling was performed simultaneously for SNPs and InDels using GATK’s v3.4 HaplotypeCaller in gVCF mode for individual samples. Genotype likelihoods were then used from the entire cohort to perform joint genotyping. Variants were separated into SNPs and InDels and variant recalibration was processed for each type of variability independently (Van der Auwera et al., 2013). Passing filter variants were annotated using Ensembl’s VEP (McLaren et al., 2010).

## Conflict of interest

The authors declare no conflict of interest.

## Acknowledgments

Jose Bras’ and Rita Guerreiro’s work is funded in part by Alzheimer’s Society. This work was supported in part by the Wellcome Trust/MRC Joint Call in Neurodegeneration award (WT089698) to the UK Parkinson’s Disease Consortium whose members are from the UCL Institute of Neurology, the University of Sheffield, and the MRC Protein Phosphorylation Unit at the University of Dundee. This work was supported in part by the Intramural Research Programs of the National Institute on Aging, Z01 AG000949-02; an anonymous charitable foundation; Alzheimer’s Research UK (ARUK); and the Intramural Research Programs of the National Institute on Aging and the National Institute of Neurological Disease and Stroke, National Institutes of Health (Department of Health and Human Services Project number, ZO1 AG000950-10). VVD and JQT’s work was supported in part by Penn Alzheimer’s Disease Core Center P30 AG-010124 and Penn Udall Center P50 NS-053488. We would also like to acknowledge the brain banks providing the tissue samples. This includes the Thomas Willis Oxford Brain Collection, the London Neurodegenerative Diseases Brain Bank, Brains for Dementia Research, the Medical Research Council and the NeuroResource. We are grateful to the Banner Sun Health Research Institute Brain and Body Donation Program of Sun City, Arizona for the provision of human biospecimens. The Brain and Body Donation Program is supported by the National Institute of Neurological Disorders and Stroke (U24 NS072026 National Brain and Tissue Resource for Parkinson’s Disease and Related Disorders), the National Institute on Aging (P30 AG19610 Arizona Alzheimer’s Disease Core Center), the Arizona Department of Health Services (contract 211002, Arizona Alzheimer’s Research Center), the Arizona Biomedical Research Commission (contracts 4001, 0011, 05-901 and 1001 to the Arizona Parkinson’s Disease Consortium) and the Michael J. Fox Foundation for Parkinson’s Research.

